# PD-L1-expressing astrocytes act as a gate-keeper for neuroinflammation in the central nervous system of mice with traumatic brain injury

**DOI:** 10.1101/2021.11.04.467368

**Authors:** Xiang Gao, Wei Li, Fahim Syed, Fang Yuan, Ping Li, Qigui Yu

**Affiliations:** Spinal Cord and Brain Injury Research Group, Department of Neurological Surgery, Stark Neurosciences Research Institute, Indiana University School of Medicine, Indianapolis, IN 46202; Department of Microbiology and Immunology, Indiana University School of Medicine, Indianapolis, IN 46202; Department of Surgery, Indiana University School of Medicine, Indianapolis, IN 46202

**Keywords:** traumatic brain injury, astrocyte, immune checkpoint, PD-L1, reverse signaling, neuroimmune response, neuroinflammation

## Abstract

**Background:** Tissue damage and cellular destruction are the major events in traumatic brain injury (TBI), which trigger sterile neuroimmune and neuroinflammatory responses in the brain. While appropriate acute and transient neuroimmune and neuroinflammatory responses facilitate the repair and adaptation of injured brain tissues, prolonged and excessive neuroimmune and neuroinflammatory responses exacerbate brain damage. The mechanisms that control the intensity and duration of neuroimmune and neuroinflammatory responses in TBI largely remain elusive.

**Methods:** We used the controlled cortical impact (CCI) model of TBI to study the role of immune checkpoints (ICPs), key regulators of immune homeostasis, in the regulation of neuroimmune and neuroinflammatory responses in the brain *in vivo*.

**Results:** We found that *de novo* expression of PD-L1, a potent inhibitory ICP, was robustly and transiently induced in reactive astrocytes, but not in microglial cells, neurons, or oligodendrocyte progenitor cells (OPCs). These PD-L1^+^ reactive astrocytes were highly enriched to form a dense zone around the TBI lesion. Blockade of PD-L1 signaling enlarged brain tissue cavity size, increased infiltration of inflammatory Ly-6C^High^ monocytes/macrophages (M/M*ϕ*) but not tissue-repairing Ly-6C^Low/^F4/80^+^ M/M*ϕ*, and worsened TBI outcomes in mice. PD-L1 gene knockout enhanced production of CCL2 that interacted with its cognate receptor CCR2 on Ly-6C^High^ M/M*ϕ* to chemotactically recruit these cells into inflammatory sites. Mechanically, PD-L1 signaling in astrocytes likely exhibits dual inhibitory activities for the prevention of excessive neuroimmune and neuroinflammatory responses to TBI through ***(1)*** the PD-1/PD-L1 axis to suppress the activity of brain-infiltrating PD-1^+^ immune cells such as PD-1^+^ T cells, and ***(2)*** PD-L1 reverse signaling to regulate the timing and intensity of astrocyte reactions to TBI.

**Conclusions:** PD-L1^+^ astrocytes act as a gatekeeper to the brain to control TBI-related neuroimmune and neuroinflammatory responses, thereby opening a novel avenue to study the role of ICP-neuroimmune axes in the pathophysiology of TBI and other neurological disorders.

## Background

Traumatic brain injury (TBI) is a major cause of death and disability worldwide, affecting more than 10 million people annually^1-3^. TBI is composed of primary and secondary injuries. Primary injury refers to the initial direct impact that causes focal and diffuse damage of the brain. Other than injury prevention, little can be done to interfere with the primary injury^4^. The secondary injury of TBI develops in minutes to hours following the initial primary injury and contributes to further damage of brain tissue^3,5,6^. In fact, secondary injury remains an important determinant of TBI outcome, as a large percentage of people with TBI develop neurological function deficits days to weeks after the event. Secondary injury is an indirect result of destructive immunological, inflammatory, neurotoxic, and biochemical cascades that are initially triggered by the primary injury. However, the exact roles of the neuroimmune and neuroinflammatory responses in the physiological and pathological processes of secondary injury are largely unknown.

Tissue damage and cellular destruction are the major events in TBI primary injury, which triggers sterile neuroimmune and neuroinflammatory responses in the absence of microbial infection^7^. Neuroimmune and neuroinflammatory responses in TBI are characterized by glial cell activation, leukocyte recruitment, and upregulation of inflammatory mediators^8,9^, and can have both beneficial and detrimental effects^9-11^. TBI damages the blood-brain barrier (BBB), enabling recruitment of circulating immune cells such as monocytes/macrophages (M/M*ϕ*), neutrophils, and activated T cells to the injured site. The accumulation of blood-born immune cells is found in the brain parenchyma of human TBI and animal models of TBI^10^. These infiltrated cells release inflammatory mediators that mobilize neural cells such as glia and immune cells to the site of injury^10^. Resident neural cells such as astrocytes and microglia are also activated and subsequently form an area of containment between injured and healthy tissues, suggesting that acute activation of resident neural cells may represent the first line of defense following TBI^10,12^. However, when resident neural cells and infiltrated immune cells become over-activated, they can induce detrimental neurotoxic effects by releasing cytotoxic substances such as pro-inflammatory cytokines (IFN-γ, IL-1β, and TNF-α)^10,13^. These inflammatory cytokines can further activate neural cells such as astrocytes to significantly affect the pathophysiology of TBI^14^. Thus, the intensity and duration of neuroimmune and neuroinflammatory responses are critical for determining their supportive or destructive effects on the central nervous system (CNS) post-TBI.

The immune responses in the brain parenchyma are tightly regulated to prevent overwhelming immune activation and subsequent inflammatory responses in this organ that has little recovery capacity^15,16^. Since immune checkpoints (ICPs) are key regulatory molecules that maintain immune homeostasis, they likely play a critical role in regulating neuroimmune and neuroinflammatory responses in TBI. ICPs consist of paired receptor-ligand molecules that exert inhibitory or stimulatory effects on immune defense, surveillance, regulation, and self-tolerance^17-21^. Under normal circumstances, ICPs regulate the breadth, magnitude, and duration of the immune responses against malignancy and infection while protecting tissues from excessive insults. In certain pathological situations such as cancer or persistent infection, the balance between ICP stimulatory and inhibitory signals is dysregulated^17-25^. Malignant cells or persistent infections can dysregulate the expression of ICPs such as CTLA-4 (cytotoxic T-lymphocyte-associated protein 4), PD-1 (programmed cell death protein 1), PD-L1 (programmed death-ligand 1), TIM-3 (T-cell immunoglobulin mucin protein 3), and LAG-3 (lymphocyte-activation gene 3) on the surface of immune cells to evade or subvert the immune response, leading to a failure or insufficiency in anti-tumor or pathogen immune attacks^22,26-28^. These important findings laid the foundation for the clinical development of ICP blockade therapies, which abrogate ICP inhibitory signals to restore and enhance the anti-tumor activity of cytotoxic T lymphocytes (CTLs)^19,20,29,30^. In fact, ICP blockade has become a revolutionary treatment for several advanced malignancies such as melanoma and lung cancer^15,20,31^.

Currently, there has been little research conducted to study the biology and clinical relevance of ICPs in the brain after TBI. We used the controlled cortical impact (CCI) model of TBI to study the role of ICPs in the regulation of neuroimmune and neuroinflammatory responses in the brain. We found that *de novo* expression of PD-L1, a key inhibitory ICP that exerts dual inhibitory signals to suppress the activity of PD-1^+^ T cells via the PD-1/PD-L1 axis and the function of PD-L1^+^ cells via a novel intrinsic reverse signaling, was robustly and specifically induced in astrocytes that were positive for GFAP (glial fibrillary acidic protein), a reactive astrocyte marker. These PD-L1^+^ astrocytes were highly enriched to form a dense zone around the TBI lesion. Blockade of PD-L1 signaling increased infiltration of inflammatory Ly-6C^High^ M/M*ϕ*, brain tissue cavity size, and motor and emotional dysfunction of TBI mice. Therefore, our study reveals that PD-L1^+^ reactive astrocytes act as a gatekeeper to control TBI-related neuroimmune and neuroinflammatory responses, thereby opening a novel avenue to investigate the role of ICP-neuroimmune axes in TBI pathophysiology. In addition, our results provide insights into the development of ICP regulators to improve TBI clinical outcomes.

## Materials and Methods

### Animals

Both male and female C57BL/6 mice at 8-12 weeks of age (Jackson Laboratories, Bar Harbor, ME) were used in all experiments as TBI occurs in both males and females. They were housed in an animal facility with a 12/12 light/dark cycle, were supplied with a standard pellet diet, and had free access to water ad libitum. All animal procedures were approved by Indiana University Institutional Animal Care and Use Committee (IACUC).

### Controlled cortical impact model of traumatic brain injury (CCI-TBI)

Mice (8-12 weeks of age) were subjected to moderate CCI-TBI or sham using sterile procedures as we previously described^32-34^. Briefly, the mice were anesthetized using a solution of 2.5% Avertin (Sigma-Aldrich, St Louis, MO) via intraperitoneal (*IP*) injection. The mouse head was fixed in the stereotaxic frame (Kopf Instruments, Tujunga, CA) using the ear bars and bite plate. A 4-mm diameter craniotomy with a midway between the bregma and lambda sutures and laterally halfway between the central suture and the temporalis muscle was performed to remove the skullcap without disruption of the underlying dura. Prior to performing CCI-TBI, the tip of the electromagnetic impactor was adjusted and kept perpendicular to the exposed cortical surface. CCI injuries were produced using an electromagnetic impactor (Impact One, Leica Biosystems, Buffalo Grove, IL). The contact velocity was set at 3.5 m/s with a cortical impact depth of 1.0 mm. This CCI setting resulted in an injury of moderate severity. After CCI-TBI, the cranial exposure was closed using sutures. Sham mice had a craniotomy without CCI impact. During surgery and recovery, the core body temperature of the animals was maintained at 36-37°C using a heating pad. All animals survived the injury and surgery.

### Administration of PD-L1 blocking or tracking antibody

TBI damages the BBB, enabling recruitment of circulating immune cells such as monocytes/macrophages (M/M*ϕ*), neutrophils, and activated T cells to the injured site. We also took advantage of the injury-induced transient breakdown of the BBB to study whether administration of PD-L1 blocking antibody (Ab) via subcutaneous (*SC*) injection could reach the injured site of the brain to exert its blocking function. PD-L1 blocking Ab (Abcam, Cambridge, MA), PD-L1-Alex 647-conjugated Ab (Abcam, Cambridge, MA), or an irrelevant IgG Ab as a control was given to mice at 24 h post-TBI at 200 μg/kg via *SC* injection. PD-L1-Alex 647-conjugated Ab was used for histological examination of Ab penetration into the brain, while PD-L1 blocking Ab was used to study the PD-L1 function in the brain.

### Collection of cerebrospinal fluid (CSF), blood, brain, and spleen

CSF was collected from mice with TBI or sham as previously reported^35^. Briefly, anesthetized mice were shaved at the back of the head above the eyes between the ears to the bottom of the neck using a razor blade for hair removal. The head of the mouse was secured using a mouse adapter. The skin and muscle were cut until the base of the skull was exposed. To collect CSF, a sharpened glass capillary (inner diameter 0.75 mm, outer diameter 1.0 mm) was placed into the capillary holder that was connected to a collecting syringe and firmly mounted on a micromanipulator. The glass capillary tip (inner diameter: 10-20 μm) was aligned to the back of the mouse head so that the sharpened point was just behind the membrane at a ∼ 30-45 ° angle. The micromanipulator was used to move the capillary tip closer to the membrane until resistance occurred. The capillary tube was gently tapped through the membrane by knocking on the micromanipulator controls under microscope monitoring. CSF was automatically drawn into the capillary tube once an opening was punctured. Approximately 5-10 μl of CSF per mouse was collected. After collection, CSF was centrifuged for 10 sec at maximal speed using a mini centrifuge, and the supernatant was stored at −80°C until use. The heat pad was used to maintain the core body temperature at 36∼37°C during CSF collection.

After the collection of CSF, mice were used for collecting blood, spleen, and brain. Blood was collected through direct cardiac blood withdrawal without an anticoagulant to obtain serum. All serum samples were stored at −80°C until use. After blood withdrawal, mice were intracardially perfused with heparinized phosphate-buffered saline (PBS) to remove remaining blood. After perfusion, the brains and spleens were dissected out and subjected to histopathological examination, immunohistochemistry (IHC) analysis, and/or preparation of single cell population for flow cytometric analysis of immune cells.

### Tissue processing and histological analysis

Tissue processing, Nissl staining, microscopy, and measurement of cortical cavity volume of the brain were performed as described in our previous report^34^. Briefly, anesthetized mice were intracardially perfused with heparinized PBS to remove blood, followed by fixation with 4% paraformaldehyde (PFA) in PBS. The brain tissues were collected and post-fixed overnight with PFA in 4°C, followed by cryoprotection in 30% sucrose for 48 h. Serial coronal sections were cut at 30 μm using a cryostat (LeicaCM 1950; Leica, Buffalo Grove, IL) and preserved at −20°C. Nissl staining was used to analyze histological changes of the brain, including lesion cavity measurement. Briefly, sections were incubated in a solution of 0.1% cresyl violet (Sigma, St Louis, MO) for 20 min. After a quick rinse in distilled water, the sections were differentiated in 95% ethanol for 3 min, followed by dehydration in 100% ethanol for 2 × 5 min. The sections were cleared in xylene for 2 × 5 min, air-dried, and mounted with DPX Mountant (Sigma-Aldrich, St Louis, MO). Slides were viewed using an upright microscopy system that was interfaced with a computer-controlled digital camera (Zeiss Axio Imager M2, Oberkochen, Germany). Images were captured and processed using AxioVision v4.8 software (Oberkochen, Germany).

To calculate the cavity volume, series of every sixth sections (30 μm thickness, 180 μm apart) covering the injured cortex were stained with cresyl violet to show the spared cortex. The boundary contours of the contralateral and ipsilateral spared cortex were drawn with a Zeiss AxioVision v4.8 software (Oberkochen, Germany). The contours enclosed volume was measured. The percent cortex of the cavity was calculated with the following formula: percentage of the cortical cavity = (contralateral cortex volume – ipsilateral spare cortex volume) / contralateral cortex volume × 100%.

### Immunohistochemistry (IHC) analysis

Brain tissue sections were incubated in blocking solution (0.1% Triton X-100, 1% bovine serum albumin, and 5% normal serum in PBS) for 1 h at room temperature, followed by overnight incubation at 4°C with primary antibodies (Abs) against mouse PD-L1, GFAP, Iba1 (ionized calcium binding adaptor molecule 1), NeuN (neuronal nuclei), NG2 (nerve/glial antigen 2), and CD45. The sections were washed 3 times with PBS, and incubated at room temperature for 2 h with fluorescent-dye conjugated secondary Abs. After staining of nuclear DNA with DAPI (4′,6-diamidino-2-phenylindole) for 2 min, the sections were washed 3 times with PBS and were mounted on the slides using Fluorescent-G mounting medium (Invitrogen, Carlsbad, CA). Final concentrations of primary Abs were used as follows: anti-GFAP (1:1000, Sigma-Aldrich, St. Louis, MO), anti-Iba1 (1:200, Abcam, Cambridge, MA), anti-NeuN (1:1000, Millipore, Billerica, MA), anti-NG2 (1:100, Abcam, Cambridge, MA), anti-PD-L1 (1:1000, Abcam, Cambridge, MA), anti-CD45 (1:200, Abcam, Cambridge, MA), or an isotype antibody as a control staining. Secondary Abs (Jackson ImmunoResearch Laboratories, West Grove, PA) were applied in a dilution of 1:1000.

### Fluorescent intensity measurement and cell counting

The intensity of PD-L1 signal was measured using ImageJ software (version 1.47; NIH, Bethesda, MD). Briefly, after staining with PD-L1, images with PD-L1-positive signaling were captured using an invert microscope (Zeiss Axio 200, Oberkochen, Germany). The contour of injured cortex (epicenter) was drawn using ImageJ software and the total PD-L1-positive signal within the area was measured. The average of total intensity from three mice was used for comparing PD-L1-positive signal in the injured cortex among the different groups. The PD-L1 and GFAP double positive reactive astrocyte was counted as described in our previous report^36^. The contours of 3 measuring areas (the block of injured cortex with 200 μm wide from different distance to the edge of cavity) were drawn. The GFAP-positive cells were quantified using optical fractionator (MBF Bioscience, Williston, VT) while PD-L1/GFAP double positive (PD-L1^+^/GFAP^+^) cells were determined using PD-L1 signal as an indicator. When PD-L1 signal showed up, the channel was switched to GFAP signal. If GFAP signal was also detected, these cells were counted as PD-L1^+^/GFAP^+^ cells.

### Preparation of single-cell population from mouse brain and spleen

After perfusion with heparinized PBS, mice were sacrificed at various time points of post-TBI or post-sham as described above. Excised brains were used to isolate single-cell population as previously described with minor modifications^37^. Briefly, harvested whole brains were cut and minced into small pieces and incubated in 10 mL Hanks’ Balanced Salt Solution (HBSS) with calcium and magnesium supplemented with 1 mg/mL collagenase B (Sigma-Aldrich, St. Louis, MO) at 37°C for 30 min with intermittent shaking. Clumps of the brain tissues were triturated and filtered through 70 μm cell strainers with the addition of 5 mL of RPMI 1640 medium supplemented with 10% fetal bovine serum (FBS, Atlanta Biologicals, San Francisco, CA). After spinning at 700 g for 10 min, the cell pellets were resuspended in 10 mL of 40% Percoll gradient (GE Healthcare, Chicago, IL) for centrifugation at 700 g for 20 min at room temperature without brake. The cell pellets were washed once with 15 mL PBS and resuspended in 1 mL of red blood cell (RBC) lysis buffer (BioLegend, San Diego, CA) for 2 min at room temperature to lyse RBCs. After centrifugation at 250 g for 10 min at 4°C, cells were harvested for flow cytometric analysis of immune cells in the brain.

Mononuclear cells were also prepared from spleens. Briefly, spleens were mechanically disrupted using the frosted ends of glass slides in 10 mL of PBS and filtered through 70 μm cell strainers, followed by centrifugation at 700 g for 5 min. Cell pellets were resuspended in 5 mL RBC lysis buffer and incubated on ice for 5 min, followed by centrifugation at 250 g for 10 min at 4°C to harvest cells for flow cytometric analysis of immune cells in the spleen.

### Flow cytometry

Single-cell suspensions of brain and spleen tissues were subjected to cell surface staining to determine the frequency and phenotype of immune cells in these tissues. Briefly, cells were first stained with the Zombie Violet™ Fixable Viability dye (FVD, BioLegend, San Diego, CA) to exclude FVD-positive dead cells and FcR blocking CD16/CD32 Ab (eBioscience, San Diego, CA), followed by staining with fluorochrome-conjugated Abs against mouse CD45, CD11b, Ly-6G, Ly-6C, F4/80, B220, CD3, CD4, CD8, PD-1, and PD-L1. All Abs were purchased from BioLegend (San Diego, CA). Appropriate isotype controls were used at the same protein concentrations as the test Abs for control staining. After washing, cells were fixed in 2% PFA and subsequently acquired using a BD LSRFortessa flow cytometer (BD Biosciences, San Jose, CA). Flow data were analyzed using FlowJo v10 software (Tree Star, San Carlos, CA). Brain-infiltrating leukocytes were defined as CD45^High^ cells that could be distinguished from CD45^Low^ glia cells. Within the CD45^High^ population, polymorphonuclear neutrophils (PMNs) were identified by Ly-6G expression, T cells as CD45^High^CD3^+^ cells, CD4 T cells as CD45^+^CD3^+^CD4^+^, CD8 T cells as CD45^+^CD3^+^CD8^+^, B cells as CD45^+^B220^+^, and M/M*ϕ* as CD45^+^CD11b^+^Ly-6G^-^. M/M*ϕ* were further divided into inflammatory Ly-6C^High^ and tissue-repairing Ly-6C^Low^F4/80^High^ subsets.

### Western blot analysis

Mice were sacrificed at 24 h, 72 h, 1 week and 2 weeks post-TBI or -sham. Entire brain tissues were collected without fixation and homogenized in the T-PER Tissue Protein Extraction Reagent (Thermo Fisher Scientific, Waltham, MA) plus 1 x Protease Inhibitor Cocktail (Sigma-Aldrich, St. Louis, MO). Tissue lysates were pulse-sonicated for 30 sec and followed by centrifugation at 14,000 x g for 15 min at 4 °C. Supernatant was harvested and subjected to the BCA Protein Assay to determine the protein concentrations. The extracted proteins were subjected to NuPAGE Novex high-performance electrophoresis (Invitrogen, Carlsbad, CA) on an 8-12% SDS-PAGE gel, and subsequently blotted onto a polyvinylidene difluoride (PVDF) membrane (Millipore, Billerica, MA). The protein bands were probed with PD-L1 Ab and detected using LI-COR Odyssey system (LI-COR Biosciences, Lincoln, NS). Anti-mouse β- tubulin Ab (Abcam, Cambridge, MA) was used as an internal control. Similar experiments were performed at least three times.

### Enzyme-linked immunosorbent assay (ELISA)

The levels of PD-L1 and CCL-2 (also known as monocyte chemoattractant protein-1 or MCP-1) in the brain lysates from mice post-TBI or post-sham were quantified using mouse PD-L1 Duoset ELISA Kit and mouse CCL-2/JE/MCP-1 Duoset ELISA Kit, both from R&D Systems (Minneapolis, MN), respectively.

### CRISPR-Cas9-mediated knockout of PD-L1 from human astrocyte cell line

PD-L1, also known as B7 homolog 1 (B7-H1) or CD274, represents the first functionally characterized ligand of the inhibitory PD-1^38^. The PD-L1 gene is highly conserved across species, suggesting its functional importance in many species^38,39^. The human PD-L1 gene is comprised of 7 exons including a short 5′ untranslated region (exon 1), a secretory signal at the amino terminus (exon 2), an IgV domain (exon 3), an IgC domain (exon 4), a transmembrane domain (exon 5), a short cytoplasmic tail (exon 6), and a long 3′ untranslated region (exon 7)^38-40^. To disrupt the PD-L1 gene at the genome level, we used a single plasmid CRISPR system with a 20-bp guide RNA (gRNA) that was aligned to its target site in exon 3 in the PD-L1 gene (5’-AGCATAGTAGCTACAGACAG-3’), designed by the GUIDES program (http://guides.sanjanalab.org/)^41^. Forward and complementary reverse oligos and exon 4 forward and complementary reverse primers containing a 20-base guide RNA (gRNA) were annealed and cloned into pSpCas9(BB)-2A-GFP (PX458)^42^, a backbone plasmid co-expressing Cas9 and EGFP (a gift from Feng Zhang; Addgene plasmid# 48138; http://n2t.net/addgene:48138; RRID: Addgene_48138), to generate a single plasmid construct co-expressing gRNA, Cas9, and EGFP. The recombinant plasmid clones, each with an insertion of a single gRNA under U6 promotor control, were screened by PCR using the universal hU6 forward primer (5’-GAGGGCCTATTTCCCATGATT-3’) and the gRNA reverse primer. The Cas9-gRNA recombinant plasmid (2.5 μg) was transfected into U87MG cells, a human astrocyte cell line (ATCC, Manassas, VA), using Lipofectamine 3000 reagents as per manufacturer’s instructions (Invitrogen, Carlsbad, CA). After 7 days of culture, PD-L1-negative cells were sorted using flow cytometry. The sorted cells were cultured and followed by an additional sorting of cells with PD-L1 knockout (KO) phenotype.

To study the regulation of PD-L1 expression and MCP-1 production, both wild-type (WT) U87MG and PD-L1 KO U87MG cells were treated with medium alone or mixed cytokines (IFN-γ, IL-1β, and TNF-α at 10 ng/mL each cytokine) for 24 h. During the last 6 h of incubation, brefeldin A (BFA, eBioscience, San Diego, CA) was added to a final concentration of 3 μM to allow intracellular accumulation of cytokines. Stimulated cells were stained with fluorochrome-conjugated anti-MCP-1 and anti-IL-6 Abs using the Cytofix/Cytoperm kit (BD Biosciences, San Jose, CA) to determine MCP-1 and IL-6 production. All experiments were repeated at least four times.

### Rotarod and elevated plus maze (EPM) tests

The rotarod test was conducted to evaluate the effects of anti-PD-L1 Ab treatment on balance and coordination of mice post-TBI versus sham as described in our previous report^43^. The rotating rod was 3 cm in diameter and divided by flanges in five compartments to allow testing of up to five mice simultaneously. The animals had to walk on the rotating rod from the lowest speed to gradually increased speed (0-30 rpm within 90 seconds) and the time until the mouse fell from the rod (latency to fall) was recorded. Mice were rested for 20 min between every two tests. A total of 3 trails were performed for each time point.

The elevated plus maze (EPM) test was performed to assess anxiety-related behavior of mice as previously described^44^. The EPM apparatus consists of a cross-shape platform with two oppositely positioned closed arms with high walls, two oppositely positioned open arms, and a small center area. The EPM apparatus was placed at 0.5 meter from the ground. The EPM test is based on the natural aversion of rodents to open spaces. Mice were allowed to start at the maze center facing one of the open arms to explore the maze for a specific amount of time. As mice freely explored the maze, their behavior was recorded by a video camera mounted above the maze. Mouse behavioral metrics including total distance traveled and time spent in each arm were analyzed using ANY-maze software (Stoelting, Wood Dale, IL). The preference for being in open arms over closed arms (expressed as moving distance and time spent in the open arms) was calculated to measure anxiety-related behavior.

### Statistical analysis

The two-sample test for two group comparison and ANOVA for multiple group comparison were used. The generalized linear mixture model (GLMM) was used to fit and test the repeatedly measured longitudinal data of Rotarod test. ANOVA test, followed by Dunnett’s test, was used to calculate differences in flow data between TBI and sham groups at different time points. Two-tailed *t* test was used to calculate differences in flow data between the anti-PD-L1 treated group and the IgG control group and differences between the wild-type and PD-L1 KO U87MG cell lines. *p*< 0.05 was considered statistically significant.

## Results

### *De novo* expression of PD-L1 was markedly and transiently induced in the brain of TBI mice

ICPs consist of paired receptor-ligand molecules that exert inhibitory or stimulatory effects on immune defense, surveillance, regulation, and self-tolerance^17-21^. The origin, regulation, and function of ICPs have been studied most extensively in the peripheral blood and peripheral tissues, but not in the CNS. Given that the neuroimmune interactions in the brain parenchyma are tightly regulated to prevent overwhelming immune activation in this organ that has little recovery capacity, we hypothesized that there were ICP axes in the CNS, which played a critical role in maintaining the neuroimmune homeostasis. To test this hypothesis, we used TBI mice to explore whether ICP expression could be induced or regulated in the brain *in vivo* in response to neuroimmune activation, as a sterile immune response develops within minutes in TBI. We performed IHC study of PD-L1 in the brain of TBI versus sham mice. As shown in Fig. 1, PD-L1 expression was barely detectable in sham mice. In TBI mice, *de novo* expression of PD-L1 was markedly and transiently induced in the brain, particularly in the cortex around the lesion area (Fig. 1A). PD-L1 expression reached the highest level on 3 days to 1-week post-TBI and declined to low or undetectable levels by 2 weeks post-TBI (Fig. 1A, 1B). PD-L1 signal was solely found in the injured side of the brain, not in the contralateral side (Fig. 1A). These IHC results were also confirmed by western blot analysis. As shown in Fig. 1C, PD-L1 expression was detected in brain tissue lysates from mice on 24 h to 1-week post-TBI and became low or undetectable by 2 weeks post-TBI. PD-L1 signal was barely detectable in brain tissue lysates from sham mice at any time point during a 2-week experiment (Fig. 1B, 1C). Thus, *de novo* expression of PD-L1 is markedly and transiently induced in response to a sterile immune response in the brain of TBI mice.

**Figure 1.**
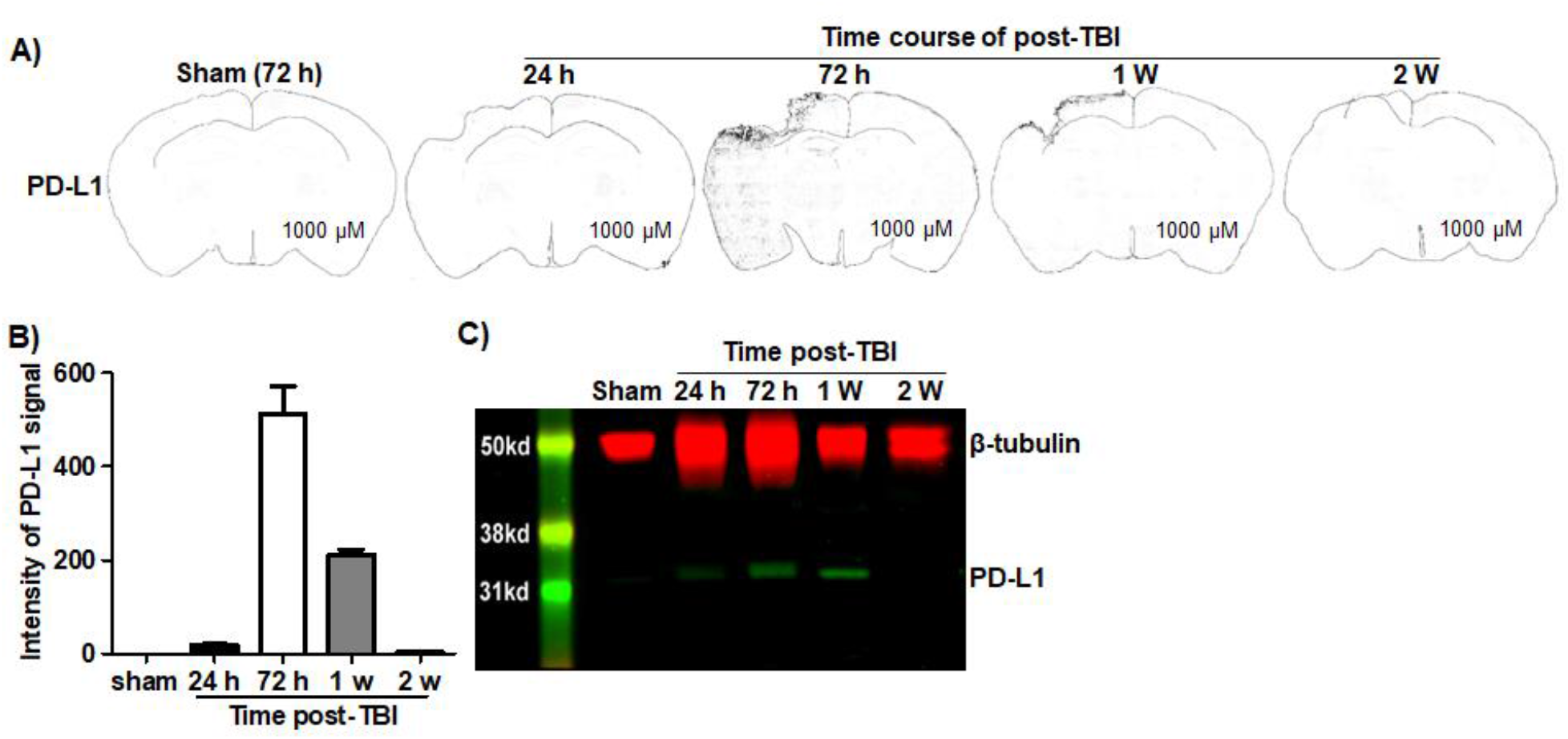
*De novo* expression of PD-L1 was markedly and transiently induced in the brain of TBI mice. After perfusion with heparinized-PBS to remove blood, the brain tissues were harvested for IHC or western blot analysis of PD-L1 expression. **A)** Coronal sections of the brain tissues from mice at different time points of post-TBI or -sham. Black dots demonstrate PD-L1-positive cells. Time course of PD-L1 expression included 72 h post-sham and 24 h, 72 h, 1 week, and 2 weeks post-TBI. **B)** Quantitation of PD-L1 expression intensity in cortex of brain coronal sections from mice at different time points of post-TBI or –sham using the ImageJ software. **C)** Western blot analysis of PD-L1 in the brain tissues of mice at different time points of post-TBI or –sham. The brain tissues of mice 72 h post-sham or 24 h to 2 weeks post-TBI were harvested for protein extraction, followed by western blot analysis of PD-L1. β-tubulin was used as an internal control. Data of IHC and western blot were obtained from 3 mice per time point.

### PD-L1 expression was transiently and specifically induced in astrocytes in the brain of TBI mice

Next, we evaluated PD-L1 expression in neural cell types in the CNS. We performed IHC analysis of PD-L1 expression in astrocytes, microglial cells, neurons, and oligodendrocyte progenitor cells (OPCs) in the brain tissues from TBI versus sham mice. As shown in Fig. 2, PD-L1 expression was transiently and exclusively induced in reactive astrocytes in the brain of TBI mice. PD-L1 expression reached the highest level on 3 days to 1-week post-TBI in reactive astrocytes, where PD-L1 and GFAP were co-localized (Fig. 2, c, c1, d and d1). The co-expression of PD-L1 and GFAP was clearly visualized in high power images (Fig. 2, f1-3 and g1-3). Thereafter, PD-L1 expression declined and became low or undetectable by 2 weeks post-TBI (Fig.2, e and e1). PD-L1 expression was undetectable in NeuN-positive neurons (Fig. 2, h, h1-3), Iba1-positve microglial cells (Fig. 2, I, i1-3), and NG2-positive OPCs (Fig. 2, j, j1-3). Thus, *de novo* expression of PD-L1 was markedly and selectively induced in astrocytes, not in other types of neural cells in the brain of TBI mice.

**Figure 2.**
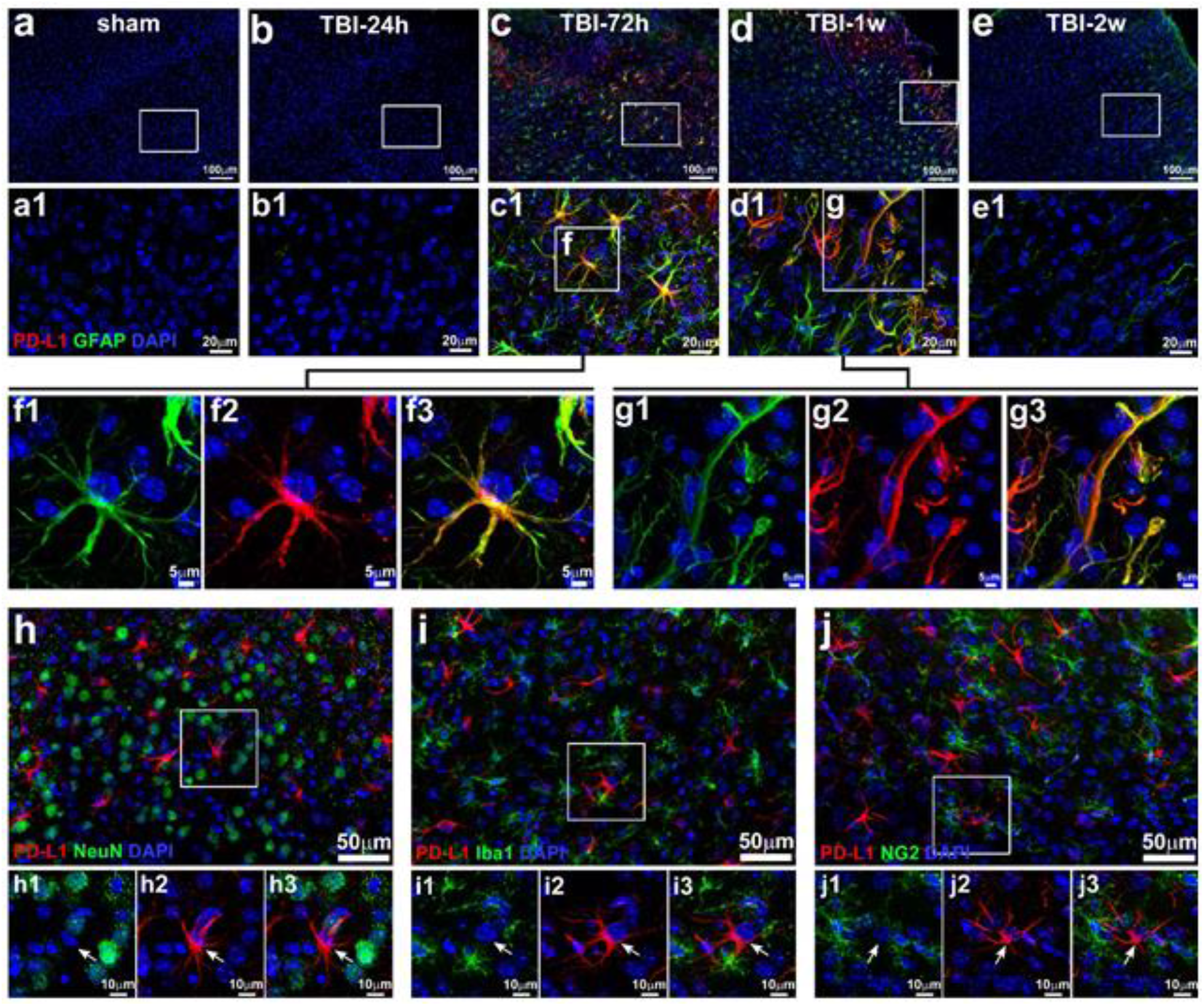
PD-L1 expression was specifically and transiently induced in astrocytes in TBI mice. Cortex sections of the brains from sham mice (a, a1) and TBI mice at different time points of post-injury, 24h (b, b1), 72h (c, c1), 1w (d, d1) and 2w (e, e1) were stained with antibodies against PD-L1 (red) and GFAP (green for astrocytes). Enlarged single cell images of the c1-f (f1-3) and d1-g (g1-3) as indicated white box showing co-localization of PD-L1 with GFAP in reactive astrocytes. h-j), low power image of IHC staining of the brain tissue sections with antibodies against PD-L1 (red) and different cell type specific marker (green) including NeuN (neurons), Iba1 (microglial cells), and NG2 (oligodendrocyte progenitor cells or OPCs). DAPI (blue) was used to counter staining nucleus in these IHC analyses. High power images showed no PD-L1 expression was detected in either Neuron (h1-3), reactive microglia (i1-3), or OPCs (j1-3).

Under normal physiological circumstances, astrocytes in the adult brain are quiescent and non-migratory cells^45^. However, astrocytes can be activated to become migratory under pathological conditions such as TBI and neuroinflammation^45^. We found that the distribution of PD-L1^+^ reactive astrocytes was changing during the course of TBI. As shown in Fig. 1A and Fig. 3, widespread PD-L1^+^ reactive astrocytes were observed in the injured cortex with highest density close to the edge of the cavity on day 3 post-TBI. Quantification of the percentage of PD-L1^+^ reactive astrocytes (PD-L1^+^GFAP^+^) in total GFAP-positive cells at different distances from the injury site showed that approximately 80% of the reactive astrocytes expressed PD-L1 within 200 μm of the injured site (Fig. 3B). The percentage of PD-L1^+^ reactive astrocytes gradually decreased with an increase in the distance away from the injured site (from ∼80%/200 μm to ∼30%/1600 μm, Fig. 3B). Strikingly, by 1-week post-TBI, PD-L1^+^ reactive astrocytes were highly enriched to form a dense zone around the TBI lesion (Fig. 2C), where the glial scar was forming. In addition, most PD-L1^+^ cells at this time exhibited scar forming astrocyte-like morphology with long and thin extensions toward the injured site (Fig. 2, d1). Notably, sporadic PD-L1^+^ astrocytes were also observed in the hippocampus of TBI mice, although the frequency was low (Fig. 2D). These results suggest that PD-L1^+^ reactive astrocytes may play a critical role in implementing a glial scar to confine TBI-triggered brain damage.

**Figure 3.**
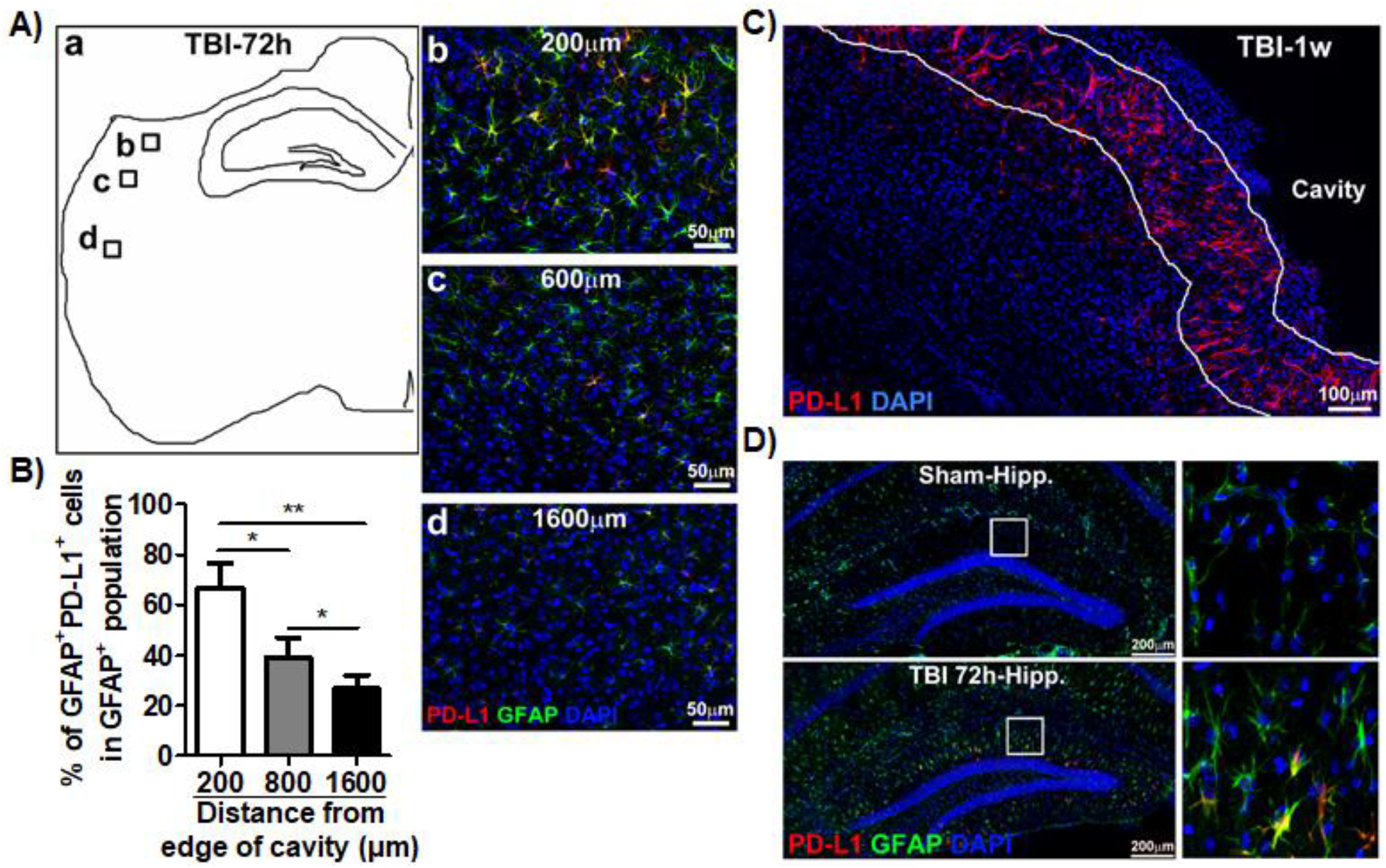
Distribution of PD-L1-positive reactive astrocytes in the brain. The brain sections from mice at different time points of post-TBI were subjected to IHC staining with DAPI (blue) and Abs against PD-L1 (red) and GFAP (green). **A)**, a. The schematic of sampling images at different positions from edge of cavity in cortex (b=200 μm, c=800 μm, and d=1600 μm). **A)**, b-d. The representative images of PD-L1 (red), GFAP (green), and DAPI (blue) labeled reactive astrocytes at different positions shown in a). **B)** Percentage of PD-L1^+^GFAP^+^ double positive astrocytes in total GFAP^+^ reactive astrocytes at different positions from edge of cavity in cortex. Data showed the gradually decrease of percentage of PD-L1^+^GFAP^+^ astrocytes away from the edge of cavity (n=3, *p*<0.05). **C)** The representative images of PD-L1^+^ astrocytes that composed a dense zone surrounding the cavity of cortex, where and when the glia scar were forming (1 week post-TBI). **D)** Sporadic PD-L1^+^ reactive astrocytes were also found in hippocampus underneath the injured cortex of TBI mice, but not sham mice.

Taken together, *de novo* expression of PD-L1 was markedly and selectively induced in astrocytes, but not in neurons, microglial cells, or OPCs, and PD-L1^+^ astrocytes were highly enriched to form a dense zone around the lesion site in TBI mice.

### Blockade of PD-L1 increased brain tissue cavity and worsen functional outcomes of TBI

Reactive astrocytes can exert biphasic functions, that is, beneficial and detrimental effects in the CNS, yet the mechanisms remain largely unclear. To clarify whether PD-L1^+^ reactive astrocytes played a beneficial or detrimental role, we applied a single dose of PD-L1 blocking Ab via *SC* injection to mice 24 h post-TBI. To verify whether PD-L1 Ab arrived at the injured site of TBI brain and was bound to reactive astrocytes, PD-L1-Alex 647-conjugated Ab was separately given via *SC* injection. After 48 h of *SC* injection, the brain tissue was collected (72 h post-TBI) and directly examined via microscopy without additional immunostaining. The result confirmed that PD-L1-Alex 647-conjugated Ab was bound to GFAP-positive reactive astrocytes in the injured cortex (suppl. Fig. 1).

To explore whether PD-L1 blockade affected the outcomes of TBI, we performed IHC to analyze brain tissue damage and used the rotarod and EPM tests to evaluate motor function and anxiety-related behavior, respectively, as described in our previous reports^46^. As shown in Fig. 4A and 4B, a larger cavity size was present in the cortex of TBI mice with PD-L1 Ab treatment when compared with IgG treatment. For the rotarod test, mice were randomly divided into 3 groups one day prior to TBI or sham. These mice were given a rotarod test to collect baseline data of mouse motor function. As shown in Fig. 4C, there was no significant difference of motor function at baseline between the groups of mice. Following treatment with IgG or PD-L1 blocking Ab, mice underwent the rotarod test 5 times per time point at days 3, 7, 14, 21, and 28 post-TBI or post-sham. The amount of time (latency to fall) that the mice stayed on the rod was recorded and anaylzed. TBI rapidly and significantly impaired the motor function of injured mice compared to sham control, as the latency on rod at day 3 post-TBI was dramatically decreased (TBI + IgG at 36.7 ± 5.9 sec versus sham at 50.3 ± 3.8 sec, *p*<0.001). The motor function of TBI mice gradually improved until there was significant difference when compared to sham controls at day 14 post-injury (TBI at 50.5 ± 5.5 sec versus sham at 56.5 ± 4.1 sec, *p*>0.05). PD-L1 blocking Ab administration not only aggravated the motor function deficit in TBI mice but also prolonged the motor function recovery (Fig. 4C). PD-L1 blockade further decreased the latency on rod at day 3 post-TBI (TBI + PD-L1 Ab at 30.3 ± 5.3 sec versus TBI + IgG at 36.7 ± 5.9 sec, *p*<0.05) (Fig. 4C). The heightened motor function impairment in TBI mice with PD-L1 blockade persisted through the entire experimental course of 28 days, remaining impaired compared to sham mice even at day 28 post-TBI (TBI + PD-L1 Ab at 44.9 ± 4.9 sec versus TBI + IgG at 52.3 ± 5.1 sec or sham at 54.2 ± 5.6 sec, *p*<0.05) (Fig. 4C).

**Figure 4.**
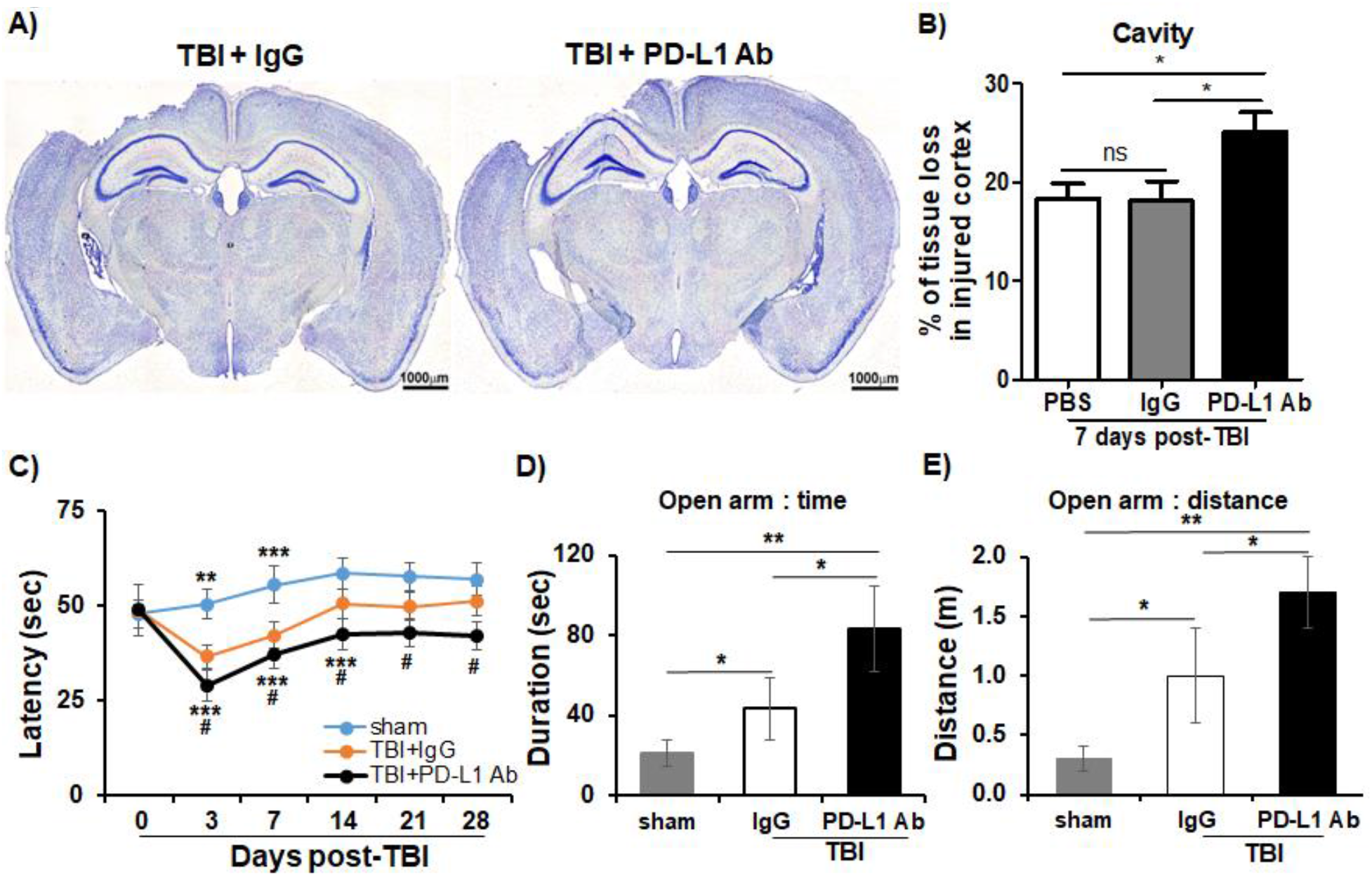
Blockade of PD-L1 increased the cavity size of injured cortex and motor and emotion dysfunction of TBI mice. After 24 h of TBI, each mouse was given a single dose (200 μg/kg) of anti-mouse PD-L1 Ab or IgG via *SC* injection. **A)** Representative images of tissue cavity in injured cortex of TBI mice treated with PD-L1 Ab or IgG using the Nissl staining. **B)** Quantitative data of tissue cavity size in injured cortex of TBI mice with PD-L1 Ab or IgG treatment (n=10/group). **C)** Rotarod test was conducted to determine the motor function. Mice received rotarod test for baseline one day before sugery and at days 3, 7, 14, 21, and 28 post-TBI or psot-sham. Data were presented as average ± SD and analyzed using the repeated measures ANOVA followed by LSD *post hoc* test. **D) and E)** The elevated plus maze (EPM) consisting of open and close arms was used to determine mouse anxious and impulsive behavior. The time (Fig. 4D) and distance (Fig. 4E) each mouse spent on the open versus close arms were recorded and calculated. Data were presented as average ± SD and analyzed using the repeated measures ANOVA. *comparison of TBI versus sham, #comparison of PD-L1 Ab versus IgG in TBI.\**p* < 0.05, \*\**p* < 0.01, \*\*\**p* < 0.001.

Since PD-L1 blockade induced prolonged impairment of motor functon, we further performed EPM tests to assess the effects of PD-L1 blockade on anxiety-related behavior, one of hippocamus-related emotional deficit, on day 28 post-TBI (27 days after PD-L1 Ab treatment). In agreement with previous findings^44^, TBI mice exhibited reduced anxiety, spending more time (TBI + IgG at 47.5 ± 11.2 sec versus sham at 24.6 ± 6.3, *p*<0.05) and traveling further (TBI + IgG at 1.1 ± 0.36 meters versus sham at 0.34 ± 0.12 meters, *p*<0.05) in the open arms of the EPM when compared to sham mice (Fig. 4D, 4E). TBI mice with PD-L1 blockade spent more time (TBI + PD-L1 Ab: 90.4 ± 17.8 sec versus TBI + IgG: 47.5 ± 11.2 sec, *p*<0.05) and traveled further (TBI + PD-L1 Ab: 1.80 ± 0.32 meters versus TBI + IgG: 1.1 ± 0.36 meters, *p*<0.05) in open arms when compared to IgG-treated mice (Fig. 4D, 4E). Of note, the movement speed of animals was not significantly different between the groups (data not shown).

Taken together, PD-L1 blockade increased the size of the brain tissue cavity, exacerbated the motor function impairment, and changed anxiety-related behavior, suggesting that PD-L1 signaling in astrocytes plays a critical role in maintaining CNS tissue integrity and functional outcomes after TBI.

### Blockade of PD-L1 prolonged the accumulation of inflammatory Ly-6C^High^, but not tissue-repairing Ly-6C^Low^/F4/80^+^ M/Mϕ in the brain of TBI mice

To study the mechanisms underlying worse outcomes of TBI due to PD-L1 blockade, we analyzed and compared immune cell infiltration to the brain versus spleen. First, we analyzed the frequency and phenotype of immune cells in the brain and spleen from heparinized PBS-perfused mice post-TBI versus post-sham at various time points. We found that the profile and frequency of immune cells were markedly altered in the brain, but less in the spleen. The percentages of neutrophils (CD45^+^Ly-6G^+^), B cells (CD3^-^B220^+^CD45^+^), and M/Mϕ (CD45^+^Ly-6G^-^ CD11b^+^) were increased in the brain, while the percentages of T cells including total T cells, CD4 T cells (CD3^+^CD4^+^), and CD8 T cells (CD3^+^CD8^+^) remained unchanged (supplemental Fig. 2). The neutrophils and B cells in the brain peaked at 24 h post-TBI, while total M/Mϕ reached their peak at 72 h post-TBI (supplemental Fig. 2). The accumulated M/Mϕ in the brain were dominated by Ly-6C^High^ M/Mϕ (CD45^+^Ly6G^-^CD11b^+^Ly-6C^High^) that were increased in a pattern similar to the total M/Mϕ. In contrast, Ly-6C^Low^F40/80^+^ M/Mϕ (CD45^+^Ly6G^-^CD11b^+^Ly-6C^Low/^F4/80^+^) decreased and reached their lowest percentages at 72 h post-TBI (supplemental Fig. 2). By day 7 post-TBI, all alterations described above returned to the levels found in sham mice (supplemental Fig. 2A). These types of immune cells in the spleen were not dramatically altered except for the Ly-6C^High^ M/Mϕ which were decreased at 24 h post-TBI (supplemental Fig. 2B). Therefore, immune cells in the CNS and peripheral lymphoid tissues exhibit different patterns of immune responses to TBI. Particularly, TBI triggers an increase of inflammatory Ly-6C^High^ M/Mϕ, but a decrease of tissue-repairing Ly-6C^Low^ M/Mϕ in the brain. The increased Ly-6C^High^ M/Mϕ are likely derived from the migration of compartments from peripheral lymphoid tissues such as spleen because the splenic Ly-6C^High^ M/Mϕ population seems to be inversely related to the brain population.

Next, we studied whether PD-L1 signaling affected the infiltration of immune cells to the brain of TBI versus sham mice. We found that PD-L1 blockade prolonged the accumulation of inflammatory Ly-6C^High^ M/M*ϕ* in the brain of TBI mice, while the percentage of tissue-repairing Ly-6C^Low^ M/M*ϕ* was decreased, albeit not to a statistical significance, in the brain of TBI mice (Fig. 5C). Other types of immune cells in the brain and spleen were not significantly affected by PD-L1 blockade. Notably, ∼25% of total T cells in the brain of TBI mice were PD-1^+^ (Fig. 5D), which likely represent a major trigger of PD-1/PD-L1 signals. IHC assays also revealed that PD-L1 blockade increased CD45^+^ lymphocytes in the injured cortex (Fig. 5E).

**Figure 5.**
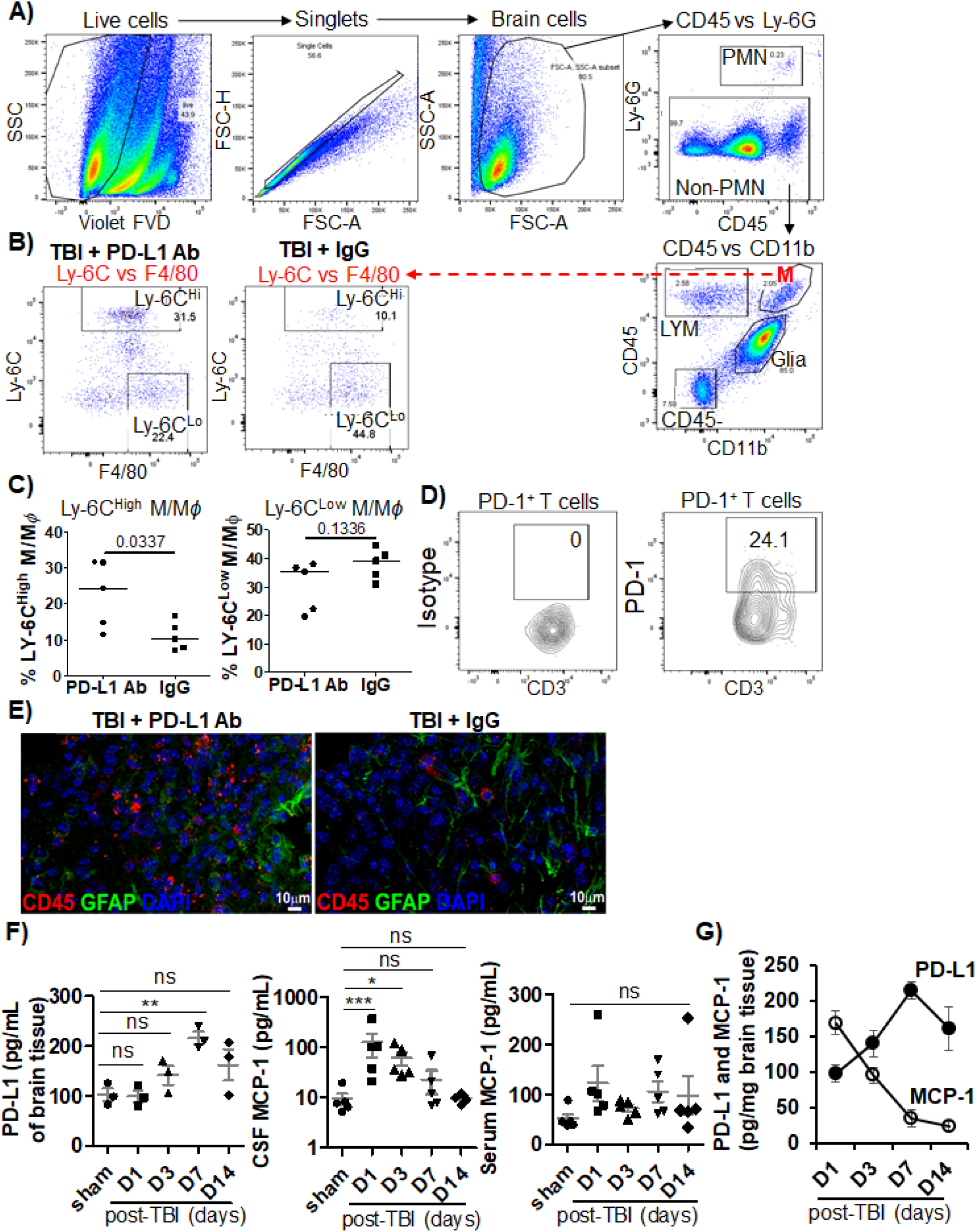
Effects of PD-L1 blockade on migration of immune cells to the brain of TBI mice. After 7 days of TBI (6 days of PD-L1 blockade), whole brain was harvested from each mouse after removal of blood cells by perfusion with heparinized-PBS, followed by preparation of brain single-cell suspension. Cells were stained with the Zombie Violet™ FVD to exclude FVD-positive dead cells, followed by incubation with CD16/CD32 Abs for blocking FcR binding, and fluorochrome-conjugated Abs against mouse CD45, CD3, CD4, CD8, CD11b, Ly-6G, Ly-6C, F4/80, B220, and PD-1. Brain-infiltrating leukocytes (LYM) were defined as CD45^High^ cells that could be distinguished from CD45^int^ glia cells. Within the CD45^High^ population, polymorphonuclear neutrophils (PMNs) were identified by Ly-6G expression, T cells as CD45^High^CD3^+^ cells, B cells as CD45^+^B220^+^, and M/M*ϕ* as CD45^+^CD11b^+^Ly-6G^-^. M/M*ϕ* were further divided into Ly-6C^High^ and Ly-6C^Low^F4/80^High^ subsets. **A)** Gating strategy for flow cytometric analysis of immune cells in the brain of TBI mice treated with PD-L1 Ab or IgG. **B)** Representative flow plots of brain cells from the gated M/Mϕ (M) showing % of Ly-6C^High^ versus Ly-6C^Low^F4/80^High^ M/Mϕ in the brain of TBI mice treated with PD-L1 Ab versus IgG. **C)** Pooled data of Ly-6C^High^ versus Ly-6C^Low^F4/80^High^ M/Mϕ in the brain of TBI mice treated with PD-L1 Ab versus IgG (n=5). \**p*<0.05. **D)** Representative flow plots showing PD-1-expressing T cells. **E)** Representative IHC of CD45^+^ lymphocytes (red) in injured cortex from mice treated with PD-L1 Ab versus IgG (n=5). **F)** ELISA results of PD-L1 in the brain lysates (n=3), MCP-1 in CSF and serum samples (n=5). **G)** The levels of PD-L1 and MCP-1 in the brain of TBI mice were present at opposite patterns of regulation. ANOVA test with Dunn’s corrections was used for comparing sham versus TBI. \**p*<0.05, \**p*<0.01, and \**p*<0.001.

Since MCP-1 (also known as CCL2) is the principle chemokine for recruiting Ly-6C^High^ M/Mϕ into inflammatory sites through binding its cognate CCR2 receptor on the cell surface^47,48^, we studied the association between levels of PD-L1 and MCP-1 in the brain of TBI mice. We used ELISA assays to quantitatively measure PD-L1 in the lysates of brain tissues from post-TBI mice, and MCP-1 in the lysates of brain tissues, CSF, and serum from mice post-sham or TBI. We found that basal levels of MCP-1 were detected in the lysates of brain tissues, CSF, and serum from mice post-sham or TBI (Fig. 5F). MCP-1 levels in the lysates of brain tissues and CSF were rapidly increased in TBI mice and reached the highest levels on day 1 post-TBI, while MCP-1 levels in the serum samples did not show any significant changes (Fig. 5F). Interestingly, we found that up-regulation of PD-L1 was associated with decline of MCP-1 in the brain of TBI mice (Fig. 5G). Thus, levels of PD-L1 and MCP-1 in the brain of TBI mice exhibit an opposite pattern of regulation, suggesting that PD-L1 may down-regulate MCP-1 production.

Taken together, PD-L1 blockade increases migration of inflammatory Ly-6C^High^ M/Mϕ to the brain of TBI mice, which is likely linked to abrogation of PD-L1-mediated suppression of MCP-1 production. Thus, PD-L1^+^ reactive astrocytes likely act as a brake on neuroinflammation in TBI.

### The effects of PD-L1 intrinsic reverse signaling in astrocytes on the production of MCP-1 (CCL2), a chemokine that recruits inflammatory Ly-6C^High^ M/M*ϕ*

PD-L1, a key inhibitory ICP, exerts dual inhibitory signals to suppress the activity of PD-1-expressing T cells via the PD-1/PD-L1 axis and the function of PD-L1^+^ cells via a novel intrinsic reverse signaling mechanism^21,30,49-65^. To test the regulatory role of PD-L1 intrinsic signaling in neuroinflammation, we used CRISPR-Cas9 technology to generate PD-L1 knockout (KO) U87MG cells, a human astrocyte cell line. As shown in Fig. 6B, wild-type (WT) U87MG cells constitutively expressed PD-L1 on the cell surface, which was not observed in primary astrocytes in the brain of mice. WT U87MG cells increased PD-L1 expression in response to stimulation with mixed cytokines (IFN-γ, TNF-α, & IL-1β) (Fig. 6B). After knockout of the PD-L1 gene at the genome level, U87MG cells completely lost PD-L1 expression in the presence or absence of stimulators (Fig. 6B).

**Figure 6.**
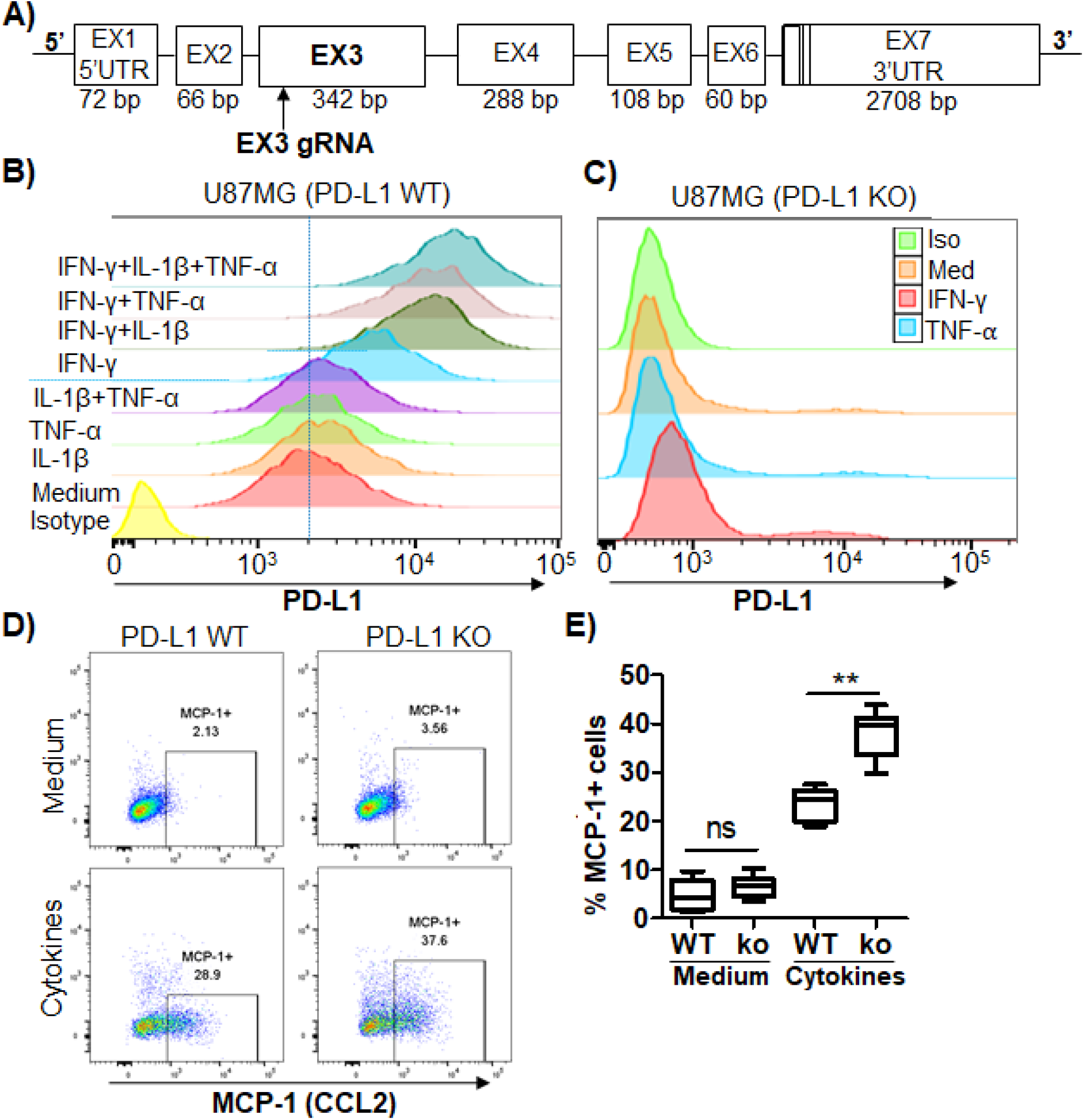
Effects of PD-L1 knockout from human astrocyte cells (U87MG) on MCP-1 (CCL2) production. **A)** Schematic representation of human PD-L1 gene structure consisting of 7 exons (EX1 – EX7, NM_014143.3) and the 20-bp guide RNA (gRNA) that was aligned to its target site in exon 3. **B)** and **C)** Representative histograms of PD-L1 expression from WT U87MG cells versus PD-L1 exon-3 KO U87MG cells. WT U87MG and PD-L1 exon-3 KO U87MG cells were treated with medium, IL-1β, TNF-α, IFN-γ, or their combinations for 24 h as indicated. Each of these cytokines was used at 10 ng/mL. Cells were subjected to surface staining of PD-L1. **D)** Representative plots of flow cytometry showing the effects of PD-L1 KO on the production of MCP-1. WT U87MG and PD-L1 exon-3 KO U87MG cells were treated with medium or mixed cytokines (IFN-γ, IL-1β, and TNF-α at 10 ng/mL each) for 24 h. Cells were subjected to intracellular staining of MCP-1 to determine MCP-1 production. **E)** Pooled data showing the effects of the effects of PD-L1 KO on the production of MCP-1 in response to stimulation with mixed cytokines. All experiments were repeated at least four times.

Next, we used PD-L1 KO versus WT U87MG cells to study the effects of PD-L1 reverse signaling in astrocytes on the production of MCP-1, a principle chemokine for recruiting Ly-6C^High^ M/Mϕ through binding to the CCR2 receptor on the cell surface^47,48^. At baseline without stimulation, both PD-L1 KO and WT U87MG cells produced little to no MCP-1 (Fig. 6D).

Stimulation with mixed cytokines (IFN-γ, TNF-α, & IL-1β) markedly increased MCP-1 production (Fig. 6D). Notably, the percentage of MCP-1-positive cells was significantly higher in PD-L1 KO than in WT U87MG cells upon stimulation with mixed cytokines (Fig. 6E). Thus, abrogation of PD-L1 reverse signaling in reactive astrocytes enhances MCP-1 production, which is likely linked to increased Ly-6C^High^ M/M*ϕ* recruitment into the brain of TBI after PD-L1 blockade.

## Discussion

This report provides evidence showing that ICPs are important and essential components of the neuroimmune system in the brain and play a critical role in the regulation of neuroimmune and neuroinflammatory responses in the brain of TBI mice. We found that *de novo* expression of PD-L1, a key inhibitory ICP that interacts with its cognate receptor PD-1 on the surface of T cells to impede T cell immunity against tumors and infectious pathogens^21,30,49-60^, was robustly induced in reactive astrocytes, but not in Iba1^+^ microglial cells, NeuN^+^ neurons, or NG2^+^ OPCs (Fig. 1 and Fig. 2). These PD-L1^+^ astrocytes were highly enriched to form a dense zone around the TBI lesion (Fig. 3). Blockade of PD-L1 signaling enlarged brain tissue cavity size, increased infiltration of inflammatory Ly-6C^High^ M/M*ϕ* but not tissue-repairing Ly-6C^Low/^F4/80^+^ M/M*ϕ*, and worsened TBI outcomes in TBI mice (Fig. 4 and Fig. 5). Mechanistically, PD-L1 signaling in astrocytes likely exhibited dual inhibitory activities in prevention of excessive neuroimmune and neuroinflammatory responses to TBI through ***(1)*** the PD-1/PD-L1 axis to suppress the activity of brain-infiltrating PD-1^+^ immune cells, especially PD-1^+^ T cells as approximately 25% of total T cells in the brain of TBI mice were PD-1^+^ (Fig. 5D), and ***(2)*** PD-L1 reverse signaling to regulate the timing and intensity of astrocyte reaction to TBI (Fig. 6). Therefore, our research suggests that PD-L1^+^ astrocytes function as a gatekeeper to the brain to control TBI-related neuroimmune and neuroinflammatory responses, thereby opening a novel avenue to study the role of ICP-neuroimmune axes in the pathophysiology of TBI and possibly other neurological disorders. Our research also provides insights into the development of ICP regulators for ameliorating post-TBI damage and promoting brain tissue repair.

In response to TBI, the brain orchestrates a complex reaction of neural and non-neural cells that interact over time to clear debris, protect viable cells, preserve function, and maintain homeostasis^66^. Astrocytes, the most abundant cells in the brain, are pivotal responders to TBI and undergo a significant change in their phenotype, gene expression, and function, resulting in the activation and proliferation, known as reactive astrocytes, astrogliosis, or astrocytosis^67^. Reactive astrocytes are heterogeneous and form scar borders to segregate damaged and inflamed tissue from adjacent neural tissue^66,68^. The most scar-forming reactive astrocytes are newly proliferated and characterized by having a large cytoplasmic mass, long branching processes, and high-level expression of intermediate filament proteins such as GFAP, vimentin, and nestin, allowing these cells to effectively communicate with and affect surrounding neural cells, non-neural cells, and neural vasculature^69^. Our studies for the first time demonstrated that reactive astrocytes exerted regulatory functions through ICP-neuroimmune axes to control inflammation and immune hyper-responses in addition to being a dense physical barrier around TBI injury. Specifically, *de novo* expression of PD-L1 was robustly and specifically induced in reactive astrocytes (Fig. 1 and Fig. 2). These PD-L1^+^ astrocytes were highly enriched to form a dense zone around the TBI lesion (Fig. 3). Abrogation of PD-L1 signaling enlarged brain tissue cavity size and worsened TBI outcomes such as motor function and anxiety in mice (Fig. 4). PD-L1^+^ astrocytes can not only suppress the activity of brain-infiltrating PD-1^+^ immune cells via the PD-1/PD-L1 axis, but also exert PD-L1 intrinsic reverse signaling to down-regulate MCP-1 (CCL2) production (Fig. 5 and Fig. 6), leading to reduction of inflammatory Ly-6C^High^ M/M*ϕ* recruitment. These results strongly imply that PD-L1^+^ astrocytes add a new layer of immune regulation and protection through dual inhibitory functions of PD-L1, providing insights into the mechanisms by which ablation of reactive astrocytes prolongs neuroinflammatory responses and intensifies the damage of TBI^70^.

Tissue damage and cellular destruction are the major events in TBI primary injury, which rapidly releases molecules of damage-associated molecular patterns (DAMPs) such as high-mobility group box protein 1 (HMGB1)^71-73^. These DAMPs bind to Toll-like receptors (TLRs) and/or the receptor for advanced glycation end products (RAGE) in neural cells such as astrocytes, microglia cells, and neurons to trigger activation of nuclear factor kappa B (NF-*κ*B) ^71-73^, leading to production of inflammatory cytokines and chemokines^7,74-81^. In fact, reactive astrocytes produce a wide range of inflammatory cytokines and chemokines including IFN-γ, IL-1β, TNF-α, TGF-β, IL-6, MCP-1 (CCL2), and CXCL10^82^. It is well established that these cytokines and chemokines are potent mediators of inflammation and inflammatory cell recruitment and play a detrimental role in secondary injury of TBI^67^. We found that CSF levels of MCP-1 (CCL2) were highly elevated in TBI mice when compared to sham mice and MCP-1 was induced in human U87MG astrocytes in response to stimulation with mixed cytokines (IFN-γ, IL-1β, and TNF-α) (Fig. 5). Abrogation of PD-L1 gene *in vitro* greatly increased MCP-1 production, indicating that PD-L1 in astrocytes exhibited its inhibitory reverse signaling to suppress MCP-1 expression. MCP-1 interacts with its cognate receptor CCR2 that is highly expressed on inflammatory Ly-6C^High^ M/M*ϕ* to chemotactically recruit these cells into inflammatory sites^47,48^. We found that inflammatory Ly-6C^High^ M/M*ϕ* were accumulated in the brain of TBI mice and further increased in the brain of TBI mice after PD-L1 blockade. Notably, tissue-repairing Ly-6C^Low^F40/80^+^ M/Mϕ that express the chemokine receptor CX3CR1^83^, but no CCR2, in the brain of TBI mice were not affected by PD-L1 blockade (Fig. 5). Based on our findings, we proposed a working model that PD-L1-expressing astrocytes act as a gate-keeper for neuroinflammation in the CNS of TBI mice. TBI-induced DAMPs and inflammatory cytokines activate astrocytes. Some cytokines such as IFN-γ can activate astrocytes and are also produced by reactive astrocytes^84^, establishing IFN-γ autocrine signaling loops to amplify astrocyte activation. In addition, infiltrated immune cells such as T cells, B cells, and M/M*ϕ* produce IFN-γ, which likely accelerates IFN-γ autocrine signals in reactive astrocytes. IFN-γ binds to the specific IFN-gamma receptors 1 and 2 (IFNGR1 and IFNGR2) to activate and recruit Janus kinases 1 and 2 (JAK1 and JAK 2) that serve as a docking site for the signaling transducer and activator of transcription 1 (STAT 1). STAT1 undergoes phosphorylation and homodimerization, and dimerized STAT1 translocates to the nucleus and subsequently binds to gamma-activated sequence (GAS) elements in the promotors of IFN-γ-regulated genes to regulate gene expression, inflammation, and cell-mediated immune responses^85^. Here, we showed that IFN-γ rapidly and potently upregulated expression of the inhibitory ICP PD-L1 in astrocytes and that PD-L1 deficiency enhanced MCP-1 production in astrocytes. Our results suggest that the PD-L1 reverse signaling acts as a brake on the activity of the IFN-γ/JAK/STAT loop in reactive astrocytes, thereby playing a key role in regulation of the timing and intensity of astrocyte reaction to TBI and in prevention of excessive neuroimmune and neuroinflammatory responses to TBI.

ICP blockade has become a revolutionary treatment for several advanced malignancies such as melanoma and lung cancer. This success has led to the investigation of ICP blockade therapies for brain metastasis of these malignancies. However, emerging evidence has demonstrated that cancer patients receiving ICP blockade therapies are at risk for developing immune-related neurological complications such as cranial polyneuropathy and autoimmune encephalitis^15,86-88^. Although these neurological complications are not very common, their incidence will likely increase as the use of ICP blockade therapies is rapidly growing. Currently, there is a lack of knowledge about ICP axes in the brain and a lack of biomarkers to predict the occurrence and severity of neurological side effects of ICP blockade therapies. Our research demonstrated that PD-L1 was remarkably induced and exhibited biological functions in the brain of TBI mice (Fig. 1 and Fig. 2), providing novel insights into the molecular mechanisms underlying immune regulation in neurological disorders.

## Conclusion

To date, more than 20 ICPs including both inhibitory and stimulatory molecules have been extensively studied in the peripheral circulation but remain elusive in the CNS. These ICPs display great diversity in their expression, regulation, and function, which is largely context or disease dependent. Thus, it is important to understand the role of ICP-neuroimmune axes in the CNS in health and disease, which will reveal important insights into the pathogenesis and treatment of CNS diseases such as TBI.

## Abbreviations

TBI: Traumatic brain injury
BBB: blood-brain barrier
M/M*ϕ*: monocytes/macrophages
CNS: central nervous system
ICP: immune checkpoint
CTLA-4: cytotoxic T-lymphocyte-associated protein 4
PD-1: programmed cell death protein 1
PD-L1: programmed death-ligand 1
TIM-3: T-cell immunoglobulin mucin protein 3
LAG-3: lymphocyte- activation gene 3
CTL: cytotoxic T lymphocyte
CCI: controlled cortical impact
GFAP: glial fibrillary acidic protein
IACUC: Institutional Animal Care and Use Committee
IP: intraperitoneal
Ab: antibody
CSF: cerebrospinal fluid
PBS: phosphate-buffered saline
IHC: immunohistochemistry
PFA: paraformaldehyde
Iba1: ionized calcium binding adaptor molecule 1
NeuN: neuronal nuclei
NG2: nerve/glial antigen 2
HBSS: Hanks’ Balanced Salt Solution
FBS: fetal bovine serum
RBC: red blood cell
FVD: fixable viability dye
PMN: polymorphonuclear neutrophil
PVDF: polyvinylidene difluoride
ELISA: enzyme-linked immunosorbent assay
B7-H1: B7 homolog 1
gRNA: guide RNA
WT: wild-type
KO: knockout
EPM: elevated plus maze
GLMM: generalized linear mixture model
OPC: oligodendrocyte progenitor cell

## Figure Legends

**Supplementary Figure 1.**
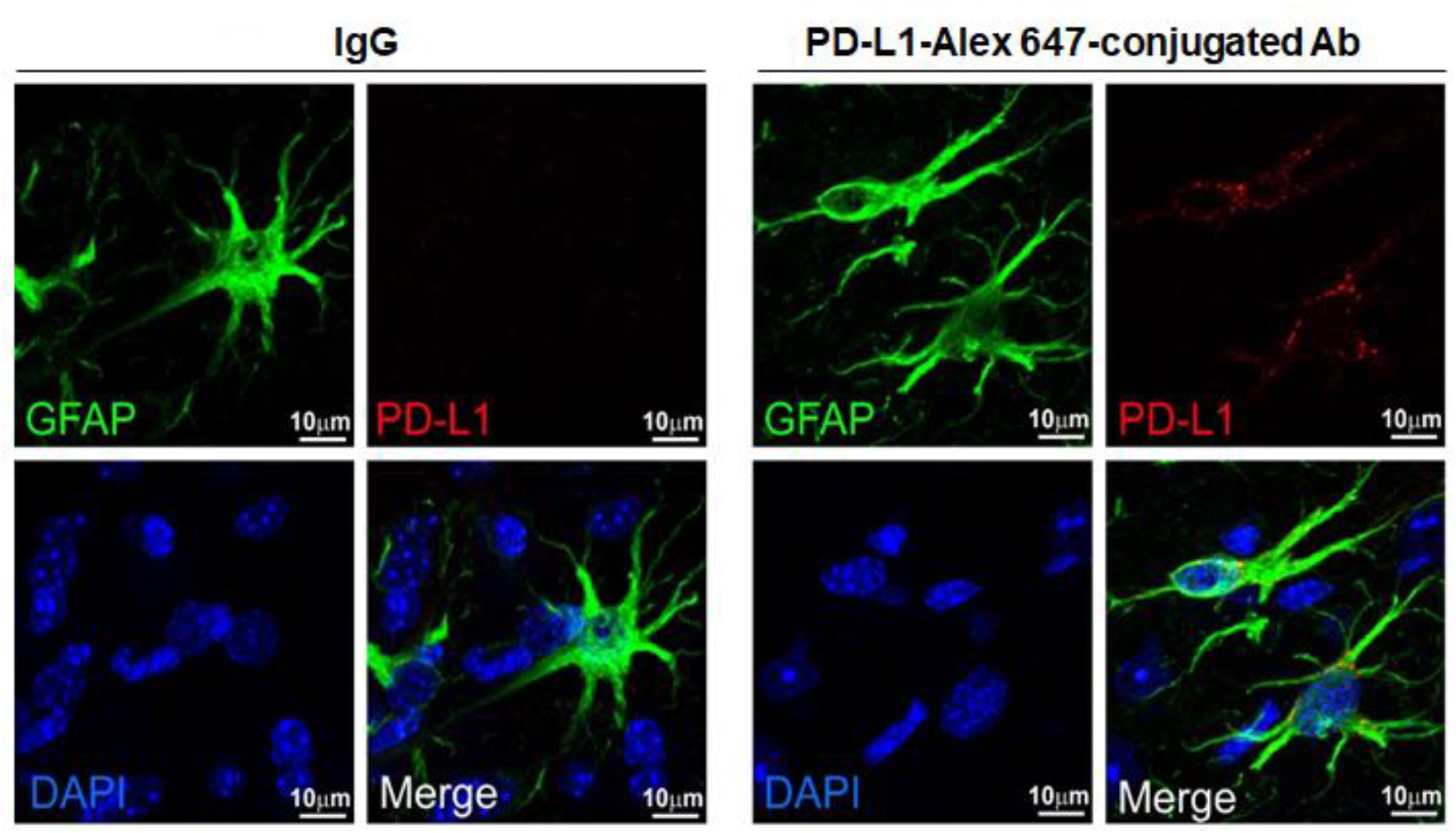
PD-L1 Ab via SC injection reached the injured site and bound to GFAP^+^ reactive astrocytes in the brain of TBI mice. Mice 24 h post-TBI were given PD-L1-Alex647-conjugated Ab (red) or IgG via SC injection. Brain tissues were harvested 48 h post-injection for IHC analysis of PD-L1 binding. Left panels, the high power images showing that there is no red signal (PD-L1) detected in the brain of TBI mice brain with IgG treatment. Right panel, the high power images showing that PD-L1-Alex647-conjugated Ab (red) reached the injured site and bound to GFAP^+^ reactive astrocytes. Similar results were obtained from 3 mice of each group.

**Supplementary Figure 2.**
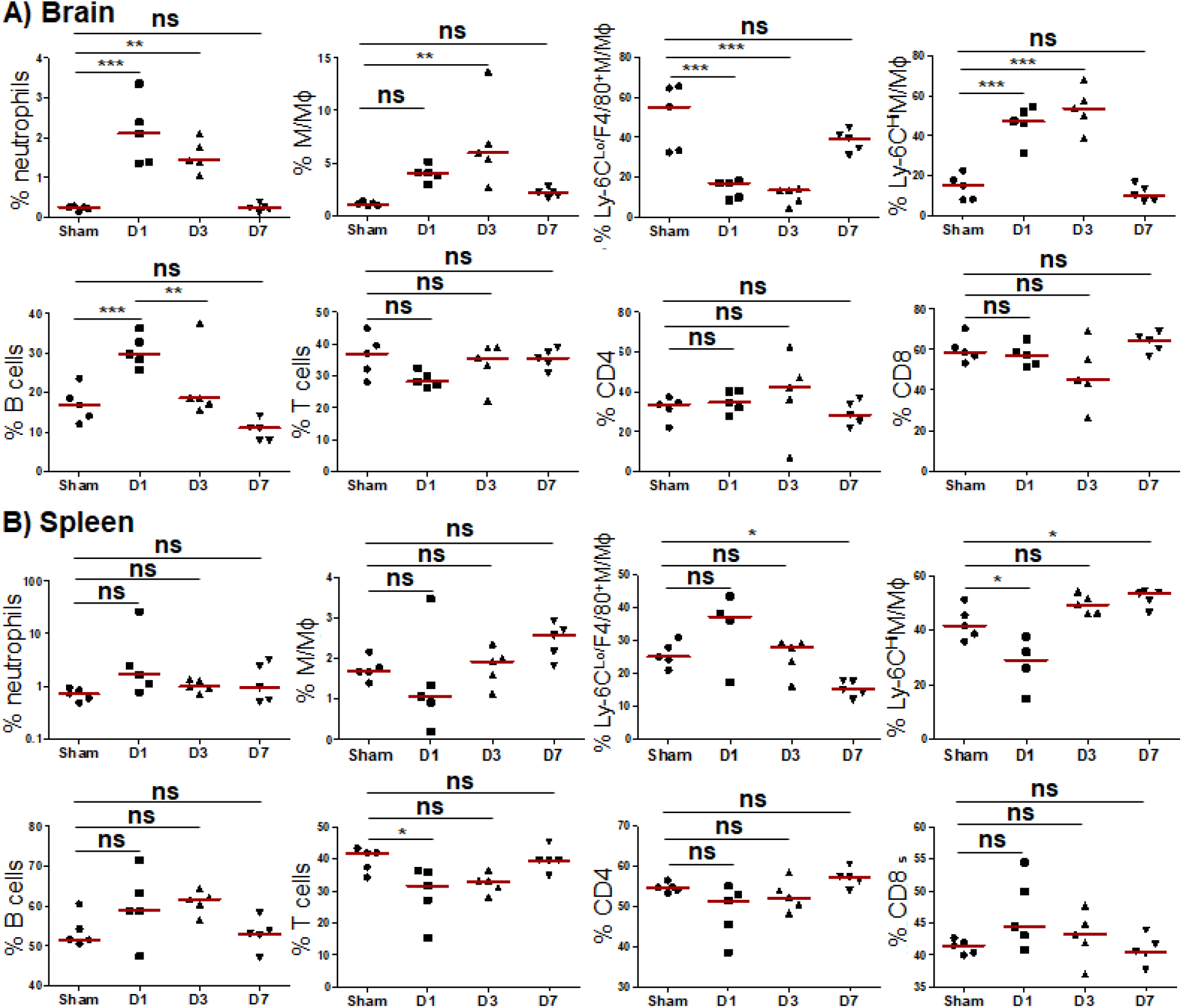
Alternations of immune cells in the brain and spleen in TBI mice. Mice (n=5/group) post-TBI or -sham were intracardially perfused with heparinized PBS to remove remaining blood. After perfusion, the brains and spleens were dissected out and subjected to preparation of single cell population for flow cytometric analysis of the frequency of immune cells including the percentages of neutrophils (CD45^+^Ly-6G^+^), B cells (CD3^-^ B220^+^CD45^+^), total M/Mϕ (CD45^+^Ly-6G^-^CD11b^+^), inflammatory Ly-6C^High^ M/Mϕ (CD45^+^Ly6G^-^ CD11b^+^Ly-6C^High^), tissue-repairing Ly-6C^Low^F40/80^+^ M/Mϕ (CD45^+^Ly6G^-^CD11b^+^Ly-6C^Low/^F4/80^+^), total T cells (CD3^+^), CD4 T cells (CD3^+^CD4^+^), and CD8 T cells (CD3^+^CD8^+^). **A)** Pooled data showing the percentage of immune cells in the brain, **B)** Pooled data showing the percentage of immune cells in the spleen. The red line represents the median. ANOVA test with Dunn’s corrections was used for comparing sham versus TBI. \**p*<0.05, \**p*<0.01, and \**p*<0.001.

## Declarations

### Ethics approval and consent to participate

Not applicable

### Consent for publication

Not applicable.

### Availability of data and materials

The data used in this study are available from the corresponding authors up on reasonable request.

### Author contributions

All authors conceived of and helped direct the study. XG, WL, FS, PL, and FY performed the experiments and analyses. All authors contributed to writing this paper.

## Acknowledgments

We would especially like to thank Dr. Jinhui Chen for his support and contributions to the conception, experimental design, data gathering, and discussion when he was in Indiana University School of Medicine.

